# The pathway of starch synthesis in *Arabidopsis thaliana* leaves

**DOI:** 10.1101/2021.01.11.426159

**Authors:** Maximilian M.F.F. Fünfgeld, Wei Wang, Hirofumi Ishihara, Stéphanie Arrivault, Regina Feil, Alison M. Smith, Mark Stitt, John E. Lunn, Totte Niittylä

**Author notes:** These authors contributed equally to this work (Maximilian M.F.F. Fünfgeld, Wei Wang). Department of Biomedical Informatics, Heidelberg University, Theodor-Kutzer-Ufer 1-3, 68167 Mannheim, Germany (Maximilian M.F.F. Fünfgeld). Corresponding authors: John Lunn and Totte Niittylä. **E-mail addresses:** Maximilian Fünfgeld, Wei Wang, Hirofumi Ishihara;, Stéphanie Arrivault, Regina Feil, Alison Smith, Mark Stitt, John Lunn, Totte Niittylä.

## Abstract

Many plants accumulate transitory starch reserves in their leaves during the day to buffer their carbohydrate supply against fluctuating light conditions, and to provide carbon and energy for survival at night. It is universally accepted that transitory starch is synthesized from ADP-glucose (ADPG) in the chloroplasts. However, the consensus that ADPG is made in the chloroplasts by ADPG pyrophosphorylase has been challenged by a controversial proposal that ADPG is made primarily in the cytosol, probably by sucrose synthase (SUS), and then imported into the chloroplasts. To resolve this long-standing controversy, we critically re-examined the experimental evidence that appears to conflict with the consensus pathway. We show that when precautions are taken to avoid artefactual changes during leaf sampling, *Arabidopsis thaliana* mutants that lack SUS activity in mesophyll cells (quadruple *sus1234*) or have no SUS activity (sextuple *sus123456*) have wild-type levels of ADPG and starch, while ADPG is 20 times lower in the *pgm* and *adg1* mutants that are blocked in the classical pathway of starch synthesis. We conclude that the ADPG needed for starch synthesis in leaves is synthesized primarily by ADPG pyrophosphorylase in the chloroplasts.

**Significance statement:** Mutant analysis shows that sucrose synthase makes no significant contribution to transitory starch synthesis in Arabidopsis leaves, resolving a 20-year old controversy about one of the most important pathways of photosynthetic metabolism.

## INTRODUCTION

Many plants accumulate starch reserves in their leaves during the day, which are remobilized at night to provide carbon and energy for survival at night (Stitt and Zeeman, 2012; Smith and Zeeman 2020). It is universally accepted that starch is synthesized from ADP-glucose (ADPG) and that in leaves starch is synthesized in the chloroplasts, primarily in mesophyll cells. However, there is disagreement about the subcellular compartmentation and pathway of ADPG synthesis. Studies on isolated chloroplasts (Heldt et al., 1977; Okita et al., 1979) and starch-deficient mutants of Arabidopsis (*Arabidopsis thaliana*; Caspar et al., 1985; Lin et al., 1988a, 1988b; Neuhaus and Stitt, 1990; Yu et al., 2000; Wang et al., 2002; Hädrich et al., 2011) led to a consensus view that ADPG is produced in the chloroplasts by ADPG pyrophosphorylase (AGPase), with its glucose 1-phosphate (Glc1P) substrate being derived from fructose 6-phosphate (Fru6P) that is withdrawn from the Calvin-Benson cycle and isomerized to glucose 6-phosphate (Glc6P) and then Glc1P by the plastidial phosphoglucomutase (PGM) and phosphoglucose isomerase (PGI), respectively. This pathway is consistent with the near absence of starch in source leaves of *pgm* and *adg1* null mutants that lack plastidial PGM or AGPase, respectively (Caspar et al. 1985; Lin et al., 1988a; 1988b). ADPG and starch synthesis are thought to be restricted to the plastids in all tissues except for the endosperm of cereals and other Poaceae, where the majority of ADPG is synthesized by a cytosolic AGPase and imported into the amyloplasts via BRITTLE-1 type ADPG transporters, probably in exchange for AMP (Denyer et al., 1996; Thorbjørnsen et al., 1996; Shannon et al., 1998; Beckles et al., 2001; Emes et al., 2003).

This consensus has been challenged by a proposal that ADPG is synthesized primarily by sucrose synthase (SUS) in the cytosol, and then imported into the chloroplasts for starch synthesis (Fig. 1B; Baroja-Fernandez et al., 2001; 2004; 2012; Munoz et al., 2005; 2006; Bahaji et al., 2011; 2013). This proposal was based on two lines of evidence. First, loss of plastidial PGM or AGPase activity in the *pgm* and the *adg1* (*aps1*) mutants, respectively, would be expected to block ADPG and starch synthesis via the consensus pathway, but leaf ADPG levels in wild-type and these mutant plants were reported not to differ significantly (Bahaji et al., 2011; Bahaji et al., 2014). Second, expression of an *Escherichia coli* adenosine diphosphate sugar pyrophosphatase (ASPP) in either the chloroplasts or the cytosol of wild-type (WT) Arabidopsis plants had effects on starch and ADPG levels that were argued to show the existence of a cytosolic pool of ADPG involved in starch synthesis. The ASPP enzyme is a phosphodiesterase that cleaves ADPG into Glc1P and AMP, but also hydrolyzes other ADP-sugars, with ADP-ribose being the preferred substrate (Moreno-Bruna et al., 2001). In plants with chloroplastic ASPP activity, ADPG levels were similar to WT but starch accumulation was decreased by about 50%. In plants with cytosolic ASPP, ADPG levels were decreased by 30% and starch accumulation was decreased by 20% (Baroja-Fernández et al., 2004). From these observations it was argued that a pool of ADPG located in the cytosol is essential for normal rates of starch synthesis. As AGPase is strictly chloroplastic in Arabidopsis leaves (Ballicora et al., 2004; Arrivault et al., 2014), Baroja-Fernandez and colleagues proposed that SUS synthesizes ADPG in the cytosol, using ADP as the glucosyl acceptor in the sucrose-cleavage reaction, followed by transfer of ADPG into the chloroplast via a hypothetical transporter in the chloroplast envelope (Baroja-Fernandez et al., 2001). However, it has not been unequivocally shown that the reported effect of cytosolic ASPP expression on starch levels is directly due to an impact on ADPG rather than a pleiotropic consequence of effects on other ADP-sugars, such as ADP-ribose.

**Figure 1.**
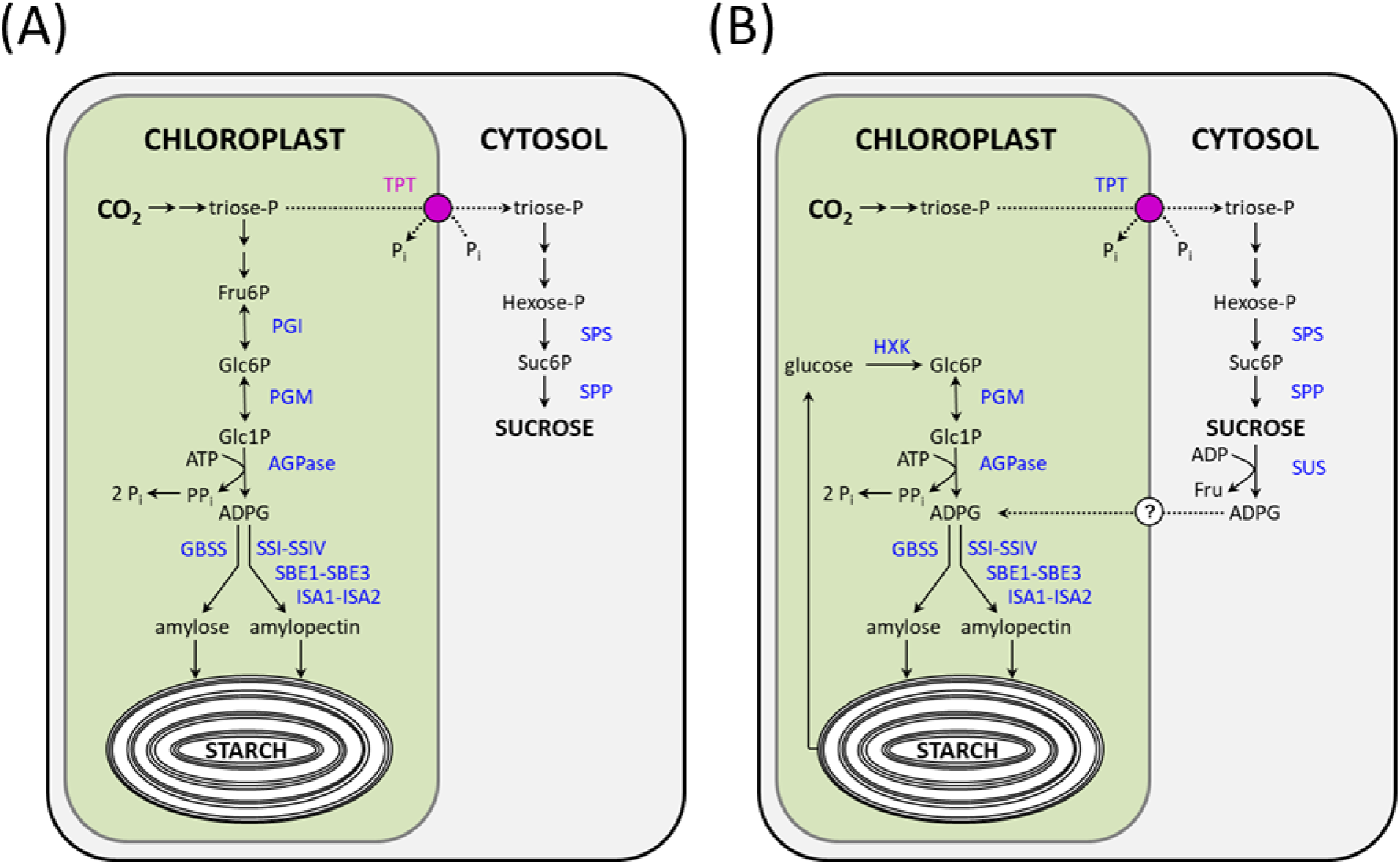
Starch synthesis in Arabidopsis leaves. (A) The classical pathway of starch synthesis (modified from a figure in Stitt et al., 2010), in which the substrate for starch synthesis, ADP-glucose (ADPG), is synthesized in the chloroplasts by ADP-glucose pyrophosphorylase (AGPase). (B) The alternative pathway proposed by Baroja-Fernandez et al. (2001) in which ADPG is primarily produced in the cytosol via sucrose synthase (SUS) and imported into the chloroplasts, via a so far unknown transporter, for starch synthesis. The model proposes that starch is turned over hydrolytically during the day producing glucose, which is phosphorylated by hexokinase (HXK) and converted back into ADPG via plastidial phosphoglucomutase (pPGM) and ADP-glucose pyrophosphorylase, for resynthesis of starch. Abbreviations: Fru6P, fructose 6-phosphate; GBSS, granule-bound starch synthase; Glc1P, glucose 1-phosphate; Glc6P, glucose 6-phosphate; ISA1, isoamylase 1; ISA2, isoamylase 2; PGI, plastidial phosphoglucose isomerase; PGM, plastidial phosphoglucomutase; SBE1-SBE3, starch branching enzymes 1-3; SPP, sucrosephosphate phosphatase; SPS, sucrose-phosphate synthase; SSI-SSIV, soluble starch synthases 1-4; Suc6P, sucrose 6’-phosphate; TPT, triose-phosphate translocator.

The proposed SUS-pathway has been critically discussed by Okita (1991) and Neuhaus et al. (2005). One of their major criticisms was that the SUS-pathway, as originally proposed, did not explain the starch-deficient phenotypes of the *pgm* and *adg1* mutants. In response to this criticism, it was proposed that a futile cycle of starch synthesis and degradation operates inside the chloroplast during the day, in parallel with synthesis and import of cytosolic ADPG. In the proposed futile cycle, starch is degraded hydrolytically to glucose, which is then phosphorylated to Glc6P by plastidial hexokinase, converted to Glc1P by plastidial PGM and then to ADPG by AGPase, for resynthesis of starch (Baroja-Fernandez et al., 2004; Munoz et al., 2006). According to this scheme, loss of either plastidial PGM or AGPase would prevent recycling of the products of starch degradation and thus prevent starch accumulation. Such futile cycling would need to operate under all conditions and throughout the photoperiod to explain the starch-deficient phenotypes of the *pgm* and *adg1* mutants. Recent ^13^C labelling studies have shown that there is some turnover of starch in Arabidopsis leaves in the light, but this is limited to certain times of day and prevalent only in long-day conditions (Fernandez et al., 2017). To test whether the proposed futile cycling of starch and glucose-salvage pathway operate in Arabidopsis leaves, Streb et al. (2009) introgressed the *pgm* mutation into null mutants for key steps in the major pathway of starch degradation in Arabidopsis leaves (Niittylä et al., 2004): glucan, water dikinase (GWD; encoded by *STARCH EXCESS 1*), phosphoglucan phosphatase (*SEX4*), isoamylase3 (*ISA3*) debranching enzyme and the maltose transporter (*MEX1*). All of the resulting double mutants were as starch-deficient as *pgm*, consistent with loss of plastidial PGM blocking the primary synthesis of starch via the consensus pathway. Even if the proposed futile cycling of starch and glucose salvage pathways were operating, the SUS-pathway does not readily explain the starch-deficient phenotype of Arabidopsis plastidial *pgi* mutants (Yu et al., 2000). Proponents of the SUS-pathway argue that plastidial *pgi* mutants are starch deficient because of a defect in isoprenoid-derived phytohormones (e.g. gibberellins) that perturb their growth and seed production, rather than a simple metabolic block in the flow of carbon from Fru6P, an intermediate of the Calvin-Benson cycle, to ADPG (Bahaji et al., 2018). The SUS-pathway also does not explain why *triose-phosphate translocator* (*tpt*) mutants, which are unable to export triose-phosphates for sucrose biosynthesis, accumulate more starch than WT plants in the light (Schneider et al., 2002). Likewise, mutants defective in the pathway of sucrose synthesis (Strand et al., 2000; Sun et al. 2011; Malinova et al., 2014) usually have elevated starch. If ADPG were synthesized primarily from sucrose via SUS, all of these mutants would be expected to have less starch than WT.

A quadruple *sus1 sus2 sus3 sus4* (*sus1234*) mutant that lacks all four of the SUS isoforms that are expressed in mesophyll cells also has WT levels of starch (Barratt et al., 2009; Yao et al., 2020), indicating that SUS is not required for starch synthesis in mesophyll cells. The main proponents of the SUS-pathway confirmed the WT starch levels in the *sus1234* mutant, but argued that the result was inconclusive as the *sus1234* mutant retains high soluble SUS activity, which they suggested had been underestimated by Barratt et al. (2009), and that Arabidopsis might have other, cryptic forms of SUS (Baroja-Fernández et al., 2012a, b).

Barratt et al. (2009) questioned whether there is sufficient SUS activity to mediate starch synthesis in Arabidopsis rosettes. This was based on a previous report of SUS activity in WT leaves (0.023 μmol min^−1^ g^−1^FW; Bieniewska et al., 2009) being lower than the rates of starch synthesis typically observed under standard laboratory conditions (approx. 0.1 μmol [Glc] min^−1^ g^−1^FW. Bieniewska et al. (2009) measured SUS activity at 25°C in the direction of sucrose synthesis, by the production of [^14^C]sucrose from UDP-[^14^C]glucose and fructose. Baroja-Fernandez et al. (2012) contended that SUS activity had been underestimated due to instability of the UDP-glucose substrate under the assay conditions. They reported an activity of 0.185 μmol min^−1^ g^−1^FW in WT leaves, measured at 37°C in the sucrose cleavage direction using an HPLC-based assay for UDP-glucose and ADPG, which would be sufficient to mediate observed rates of starch synthesis. The basis for these claims was in turn questioned by studies showing that UDP-glucose is not as unstable at moderately alkaline pH as claimed by Baroja-Fernandez et al., 2012 (Hill et al., 2017). In an independent study, Fallahi et al. (2008) measured SUS activity enzymatically at 25°C in the sucrose cleavage direction, reporting activities of 2.9 μmol min^−1^ mg^−1^ protein and 0.2 μmol min^−1^ mg^−1^ protein in rosette leaves of non-flowering and flowering plants, respectively. Assuming a typical value of 10 mg protein g^−1^FW in Arabidopsis leaves (Ishihara et al., 2015), these values translate to approx. 29 and 2 μmol min^−1^ g^−1^FW, respectively, which would be ample to support typically observed rates of starch synthesis.

In addition to the genetic evidence favouring the consensus pathway, there are biochemical data that conflict with the proposed SUS-pathway. When WT Arabidopsis rosettes were labelled with ^13^CO_2_ under steady state photosynthetic conditions, the pool of ADPG was very quickly labelled, with similar kinetics to Calvin-Benson cycle intermediates and more rapidly than sucrose, the putative precursor of ADPG synthesis in the SUS-pathway (Szecowka et al., 2013). These labelling patterns are consistent with ADPG being synthesized directly from intermediates of the Calvin-Benson cycle in the chloroplasts, rather than via sucrose in the cytosol.

Transitory starch is a major end product of photosynthesis, therefore, uncertainty about its synthesis is a major obstacle to understanding the regulation of photosynthetic carbon metabolism and how the fixed carbon is allocated between storage and export for growth. To resolve the long-standing controversy surrounding the pathway of starch synthesis in leaves, we critically examine the main lines of evidence for synthesis of ADPG via SUS or other, unknown, reactions in the cytosol. Furthermore, we generated sextuple Arabidopsis *sus1 sus2 sus3 sus4 sus5 sus6* (*sus123456*) mutants that completely lack SUS activity, and report that these mutants have near wild-type levels of ADPG and starch, and only mild growth phenotypes under standard laboratory growth conditions.

## RESULTS

### Optimization of ADPG measurement in Arabidopsis leaves

High-performance liquid chromatography with UV detection at 260 nm (HPLC-UV) has been widely used to assay ADPG, including all but the most recent study (Bahaji et al., 2014) claiming support for the SUS-pathway. However, plant extracts contain many other types of nucleotide that absorb in the UV range (Dawson et al. 1986), and it is difficult to achieve baseline separation of ADPG from other UV-absorbing compounds, potentially leading to overestimation of ADPG. HPLC coupled to tandem mass spectrometry (LC-MS/MS) offers much greater specificity and sensitivity than HPLC-UV (Lunn et al., 2006), and has become the method of choice for assaying ADPG.

The levels of ADPG in Arabidopsis leaves are generally low (<5 nmol g^−1^FW), with an estimated turnover time of <1 s in illuminated leaves (Arrivault et al., 2009), suggesting that leaf sampling in the light must be done with great care to avoid artefactual decreases in ADPG content below the steady state level before the tissue is quenched. To assess the sensitivity of ADPG to shading, rosettes were harvested from WT, *pgm* and *adg1* Arabidopsis plants in the light by either pouring liquid nitrogen directly onto the plants, or by cutting the hypocotyl from below and then quickly transferring the cut rosette into liquid nitrogen while in the growth chamber, taking care not to shade the leaves at any time during the procedure. In parallel, rosettes were harvested from plants that had been deliberately shaded by transfer from 160 μE m^−2^ s^−1^ to <10 μE m^−2^ s^−1^ for 2-3 s. Shading of WT plants for just a few seconds led to a large (94%) fall in rosette ADPG levels (measured by LC-MS/MS) compared to rosettes that had been carefully harvested in the light without shading (Fig. 2A). The *pgm* mutant contained much lower levels of ADPG than WT plants, but there was no significant effect of shading (Fig. 2B), while the *adg1* mutant had even lower levels than *pgm* which were decreased even further by shading (Fig. 2C). These results demonstrate the sensitivity of ADPG in WT plants to artefactual changes due to shading during leaf sampling, hence the need for extremely rapid quenching under full illumination. Pouring liquid nitrogen directly onto the plants is inconvenient and potentially hazardous, so for subsequent experiments we cut the hypocotyl from below and immediately transferred the cut rosette into liquid nitrogen within the growth chamber.

**Figure 2.**
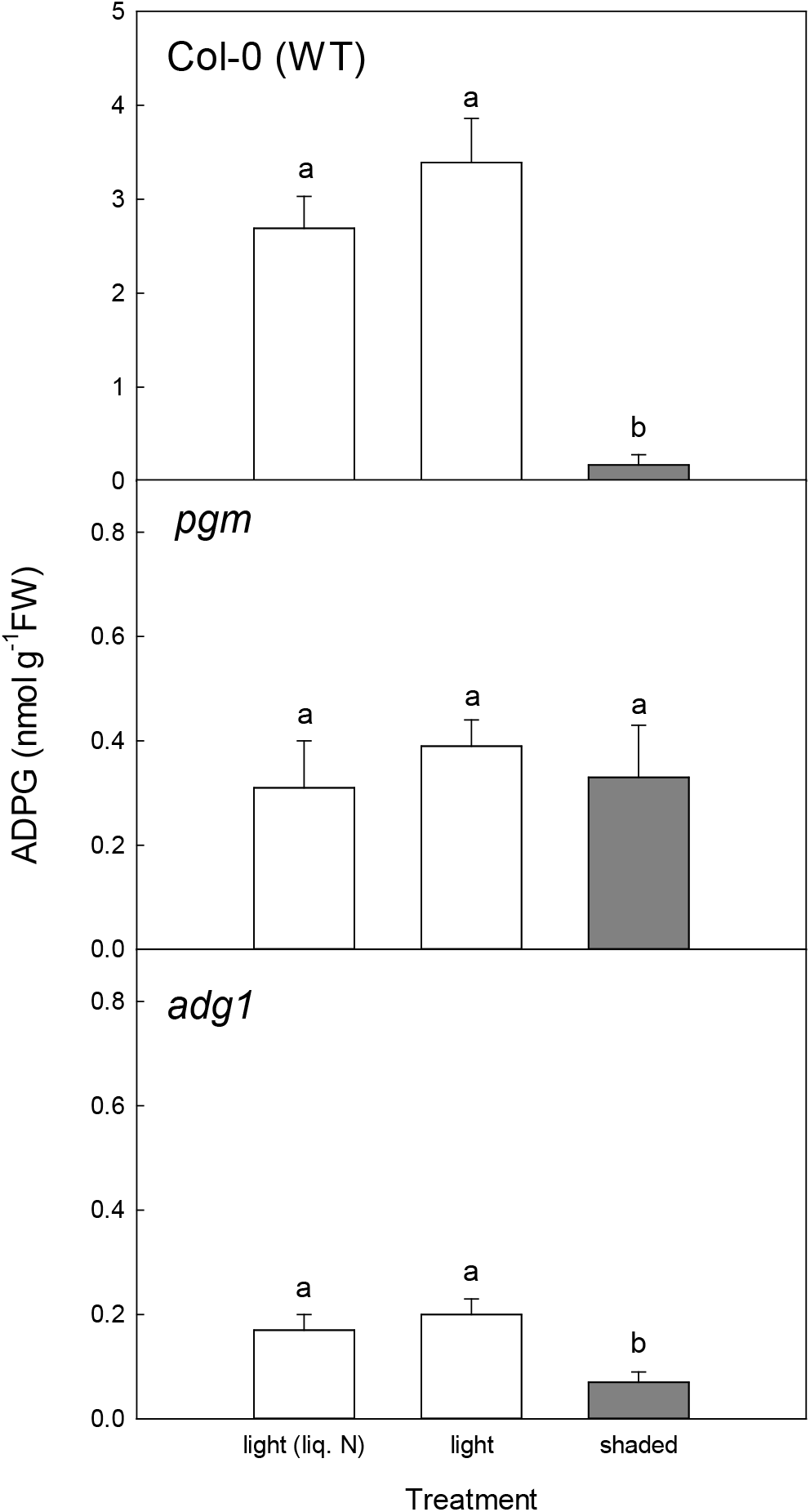
Optimization of Arabidopsis leaf harvesting measurement of ADPG. Rosettes were harvested from vegetative wild-type Col-0 plants and from starch deficient *pgm* and *adg1* mutant in the light (white bars) by pouring liquid nitrogen (liq. N) directly on to the rosette or by cutting the hypocotyl from below and transferring the cut rosette into liquid nitrogen, taking care to avoid all shading. In parallel, other plants were deliberately shaded (grey bars) for 2-3 sec before cutting and snap-freezing the rosettes. ADPG was measured by LC-MS/MS. Data are mean ± S.D. of six independently pooled batches of five rosettes (*n*=6). Letters indicate significant differences (*P*<0.05) according to a one-way ANOVA (Holm-Sidak test).

### Comparison of ADPG levels in wild-type and mutant Arabidopsis plants

WT, *pgm*, *adg1* and *sus1234* plants were grown in long-day (16-h photoperiod) conditions, and rosettes were harvested at the end of the night and at intervals during the day for metabolite analysis. There were no significant differences in Glc6P and Glc1P between WT and *sus1234* plants (Fig. 3A-B; Supplementary Table S1). In the *pgm* and *adg1* mutants, Glc6P was lower than WT in the dark, but increased to a greater extent upon illumination and was significantly higher than WT from about ZT4 onwards (Fig. 3A). Glc1P was lower than WT in the *pgm* mutant but higher than WT in *adg1* (Fig. 3B). ADPG levels were extremely low (<0.005 nmol g^−1^FW) in all genotypes in the dark (Fig. 3C). Upon illumination, ADPG increased rapidly in WT and *sus1234* plants, reaching a similar level (0.51-0.71 nmol g^−1^FW) in the two genotypes and remaining fairly constant throughout the day with no significant differences between WT and *sus1234* plants (Fig. 3C). In contrast, ADPG rose only slightly upon illumination of *pgm* and *adg1*, reaching levels of 0.03 nmol g^−1^FW and 0.01-0.02 nmol g^−1^FW, respectively (Fig. 3C). The *sus1234* plants accumulated the same amounts of starch as WT plants, whereas at ZT12 the starch contents of the *pgm* and *adg1* mutants were <1 % of WT (Fig. 3D).

**Figure 3.**
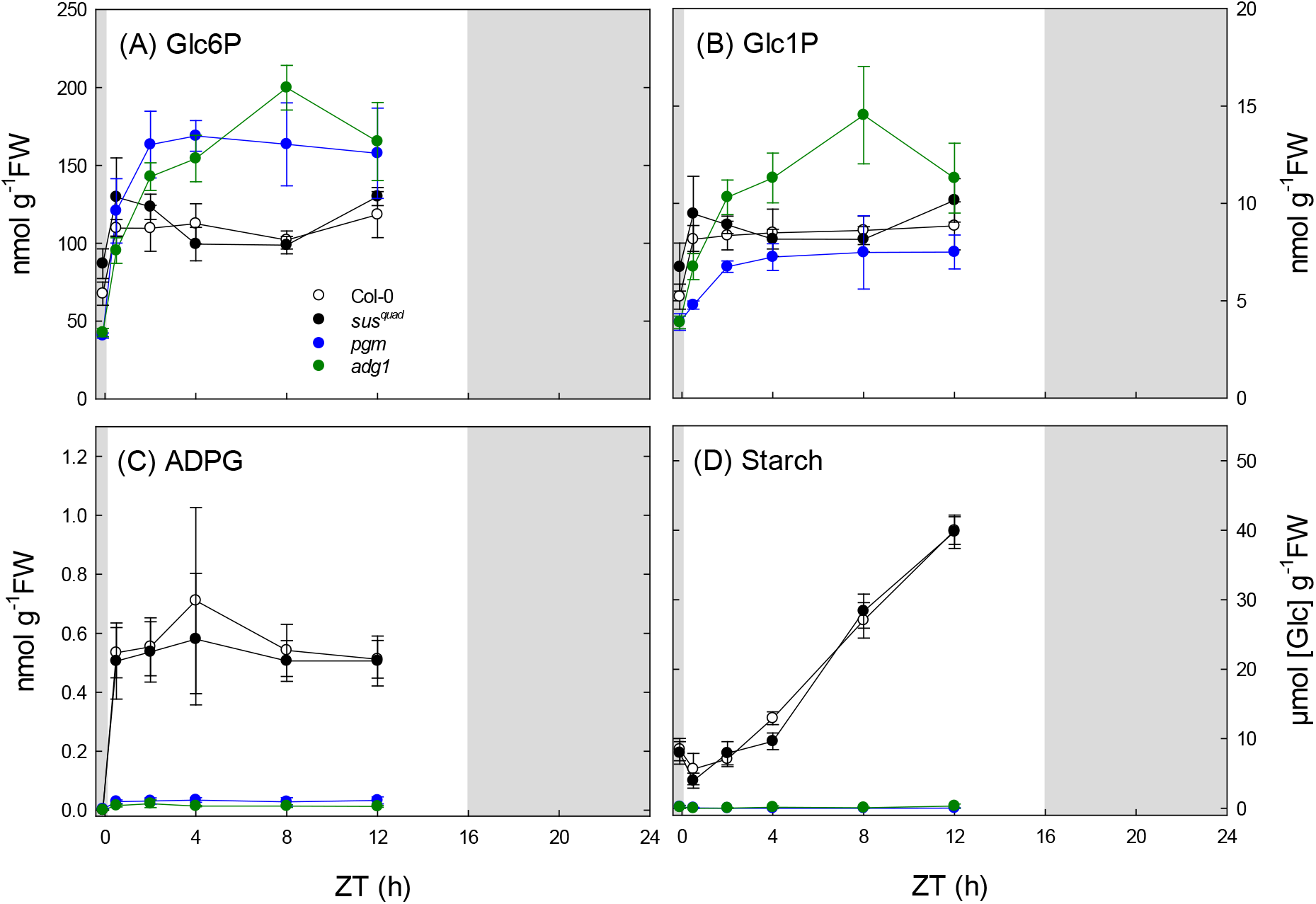
Metabolite levels in wild-type and mutant Arabidopsis plants. Wild-type Col-0, the *sus1234* mutant (*sus^quad^*) and two starch-deficient mutants (*pgm* and *adg1*) were grown in long-day conditions (16 h photoperiod). At 25 days after germination, rosettes were harvested just before dawn (ZT-0.2) and at intervals from ZT0.5 to ZT12 for measurement of: (A) glucose 6-phosphate (Glc6P), (B) glucose 1-phosphate (Glc1P), (C) ADP-glucose (ADPG) and (D) starch. Data are mean ± S.D. of four independently pooled batches of five rosettes (*n*=4). *P*-values for all genotype x genotype comparisons are shown in Supplemental Table S1.

Based on localization of the SUS1-SUS4 isoforms (Barratt et al., 2009; Yao et al., 2020), the quadruple *sus1234* mutant is expected to lack SUS activity in the mesophyll cells where transitory starch is synthesized (Smith et al., 2012). The residual SUS activity in this mutant is disputed, with Barratt et al. (2009) reporting that soluble SUS activities in roots and stems of the quadruple *sus1234* mutant were less than 2% of those in WT plants, whereas Baroja-Fernández et al. (2012a) reported soluble SUS activities of 98% and 90% of WT in leaves and stems of the *sus1234* mutant. To resolve these discrepancies, we generated two independent sextuple *sus123456* mutants by CRISPR/Cas9 mediated gene editing of the *SUS5* and *SUS6* genes in the *sus1234* mutant background, and compared the SUS activities in siliques of WT, *sus1234* and *sus123456* plants.

The CRISPR/Cas9 generated mutations in the *SUS5* (*sus5-1* and *sus5-2*) and *SUS6* (*sus6*) genes gave rise to premature stop codons (Supplementary Fig. S1) within the respective regions coding for the catalytic glucosyltransferase domains of the enzymes. The zygosity of the *sus5* and *sus6* alleles was tested by genomic PCR using gene specific primers followed by restriction enzyme digest of the PCR products (Supplementary Fig. S2). The presence of homozygous T-DNA insertions in the *SUS1, SUS2, SUS3* and *SUS4* genes was confirmed in both of the gene-edited lines (Supplementary Fig. S2), establishing that the homozygous sextuple mutants lacked functional copies of all six Arabidopsis *SUS* genes.

To confirm loss of SUS activity, we measured activity in developing siliques of WT, *sus1234*, and the two sextuple *sus123456* mutants. Developing siliques were chosen because all six *SUS* genes are expressed in siliques and/seeds (Supplementary Fig. S3), and SUS activity is high and readily quantified in these tissues (Fallahi et al., 2008). Activity was determined as the UDP-dependent production of UDP-glucose from sucrose, with UDP-glucose being measured enzymatically, as described by Dancer and ap Rees (1989). The soluble SUS activity in WT Col-0 siliques was 86.8 ± 16.7 nmol min^−1^ g^−1^FW. Activities were much lower or undetectable in the *sus1234* (1.6 ± 1.9 nmol min^−1^ g^−1^FW; <2% of WT), *sus12345^1^6* (−0.2 ± 2.9, nmol min^−1^ g^−1^FW, i.e., no detectable activity) and *sus12345^2^6* (0.2 ± 2.0 nmol min^−1^ g^−1^FW; <0.2% of WT) mutants (Fig. 4A). The residual activity in the latter was within the range of values obtained with a boiled extract of Col-0 siliques (0.6 ± 1.2 nmol min^−1^g^−1^FW) indicating it may just reflect noise at the detection limit of the assay (Fig. 4A). We conclude that the sextuple *sus123456* mutants have no detectable SUS activity. The *sus123456* mutants showed near wild-type growth under standard laboratory conditions (Fig. 4B-C).

**Figure 4.**
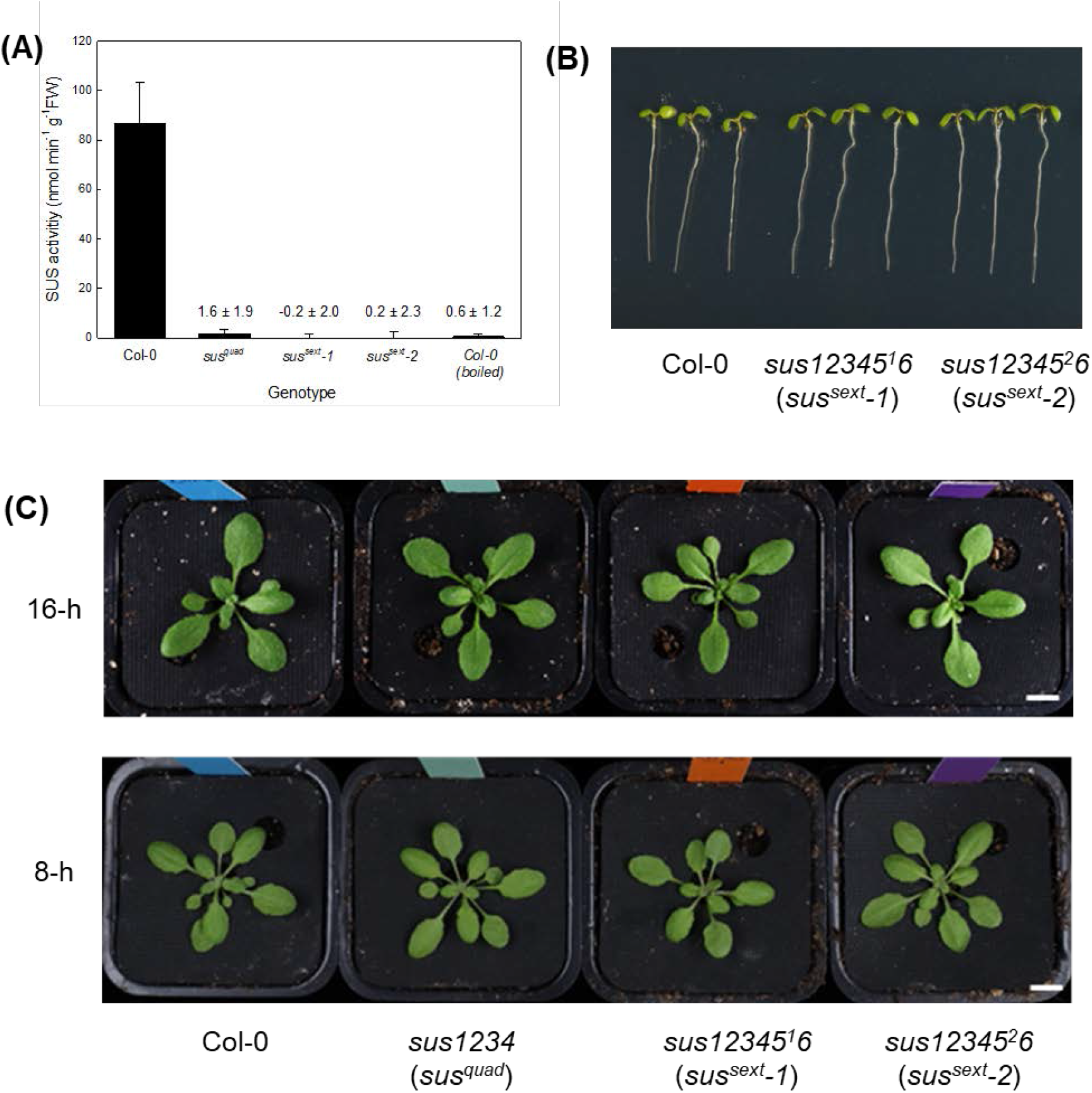
Sucrose synthase activity and morphology of Arabidopsis wild type and *sus* mutant plants. (A) Sucrose synthase activity of developing siliques, measured as the UDP-dependent production of UDP-glucose from sucrose. Data are mean ± S.D. of six independently pooled batches of five rosettes (*n*=6). (B) Seedling morphology at 5 days after germination on ½ MS medium. Bar = 5 mm. (C) Rosette morphology of plants grown in short-day (8-h photoperiod) conditions for 4 weeks. Bar = 2 cm. Abbreviations: Col-0, wild type Columbia-0; *sus^quad^, sus1234; sus^sext^-1, sus12345^1^6; sus^sext^-2, sus12345^2^6*

WT Col-0, *sus1234* and *sus12345^1^6* plants were grown under long-day (16-h photoperiod) conditions and rosettes were harvested just before dawn and at intervals during the day for metabolite analysis. In WT Col-0 plants, Glc6P, Glc1P and ADPG increased upon illumination (Fig. 5A-C), as observed previously (Fig. 3), and the plants accumulated starch in a linear manner (Fig. 5D). The phosphorylated intermediates and starch showed essentially identical levels and diel patterns in the *sus1234* and *sus12345^1^6* plants (Fig. 5), with no significant differences from WT Col-0 plants (Supplementary Table S2). Likewise, there were no significant differences between the mutant and WT Col-0 plants in the levels of soluble sugars (sucrose, glucose and fructose) or intermediates of sucrose synthesis (Fru6P, UDP-glucose, dihydroxyacetone-phosphate; Supplemental Fig. S4 and Supplemental Table S2).

**Figure 5.**
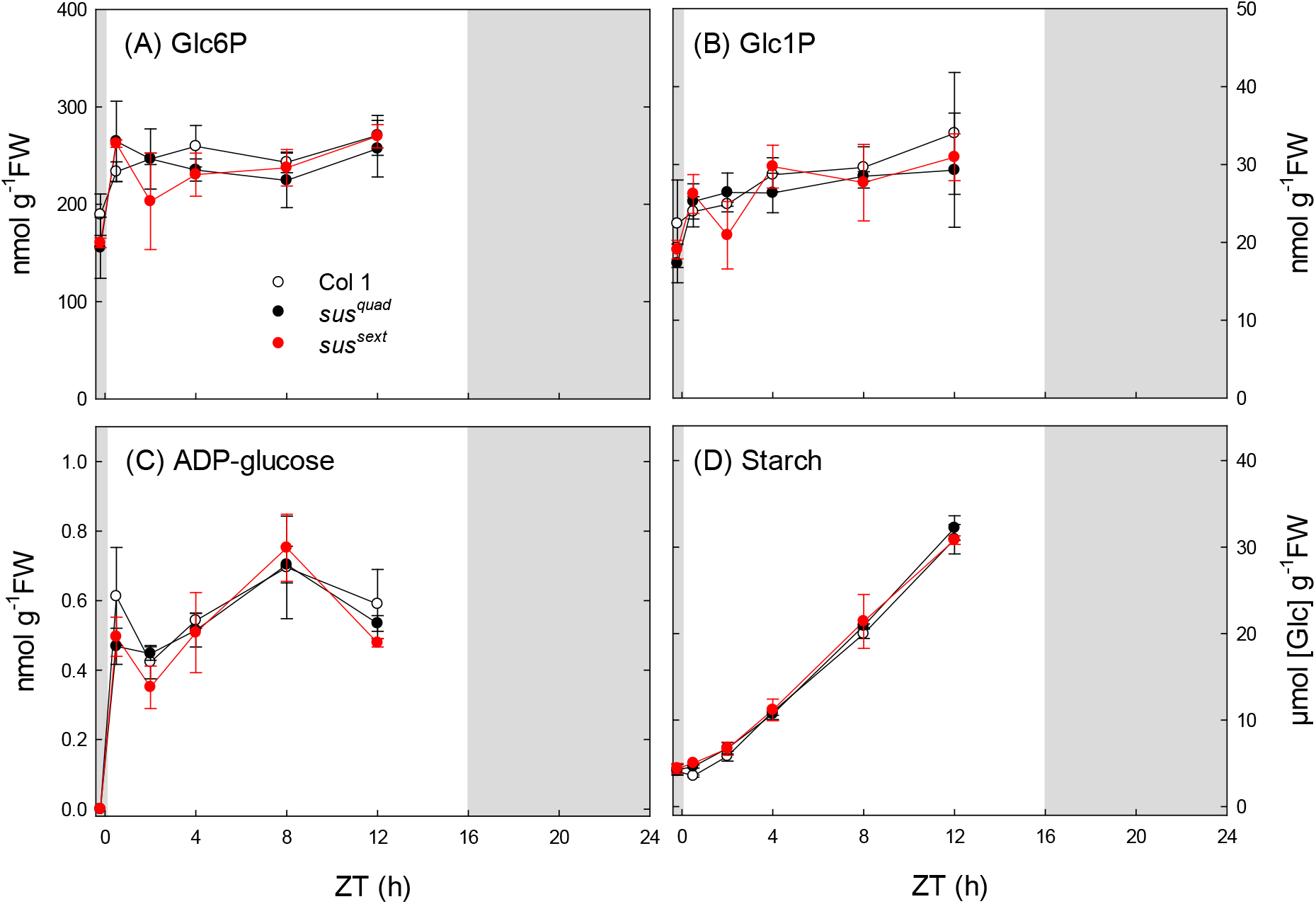
Metabolite levels in wild-type Arabidopsis and *sus* mutants. Wild-type Col-0, and the *sus1234* mutant (*sus^quad^*) and *sus12345^1^6* (*sus^sext^-1*) mutants were grown in long-day conditions (16 h photoperiod). At 25 days after germination, rosettes were harvested just before dawn (ZT-0.2) and at intervals from ZT0.5 to ZT12 for measurement of (A) glucose 6-phosphate (Glc6P), (B) glucose 1-phosphate (Glc1P), (C) ADP-glucose (ADPG) and (D) starch. Data are mean ± S.D. from three independently pooled batches of five rosettes (*n*=3). *P*-values for all genotype x genotype comparisons are shown in Supplemental Table S2.

## DISCUSSION

According to the consensus pathway, the starch-deficient *pgm* and *adg1* mutants should have little or no ADPG due to loss of essential enzymes for its synthesis in the chloroplasts. Therefore, reports of near-WT levels of ADPG in these two mutants (Bahaji et al., 2011, 2014) appeared to be inconsistent with expectations. To investigate the reliability of these reports, we established a robust method for harvesting Arabidopsis rosettes in the light that maintained *in-vivo* levels of metabolites like ADPG that are present in very small amounts and turn over very quickly (half time <1s; Arrivault et al., 2009), and used LC-MS/MS (Lunn et al., 2006; Arrivault et al., 2009) to assay ADPG with greater sensitivity and specificity than the HPLC-UV method that was used for most previous studies. We found that shading of the leaves of WT plants for just a few seconds leads to a rapid decrease in measurable ADPG content, down to levels that resemble those found in the starch-deficient *pgm* and *adg1* mutants. Consequently, unless extreme care is taken during sample collection, ADPG levels are likely to be under-estimated and therefore unrepresentative of the levels during steady state photosynthesis. This issue may well have affected values presented in earlier papers, including some of our own.

Using our optimised methods, we found ADPG levels in illuminated leaves of the *pgm* and *adg1* mutants were <6% and <4% of WT, respectively, but the same as WT in the *sus1234* mutant (Fig. 3C). In contrast, and using a similar LC-MS based assay method as us, Bahaji et al. (2014) reported that ADPG levels were the same in wild type leaves and leaves of the *pgm* and *aps1* mutants (*aps1*, like *adg1*, lacks a subunit of AGPase and is nearly starchless). In earlier work that employed the HPLC-UV assay for ADPG, these researchers reported that *adg1* leaves also had ADPG levels comparable with those of wild-type leaves (Bahaji et al., 2011). One possible explanation for this discrepancy between our results and those of Bahaji and colleagues is that their plants were inadvertently shaded during harvest, potentially leading to large losses of ADPG prior to quenching. Consistent with this idea is the fact that their values for ADPG in WT leaves are much lower than our values (Bahaji et al., 2014, reported levels of 0.13 nmol g^−1^FW compared with around 3 nmol g^−1^FW in our experiments shown in Fig. 2A). This low value for WT leaves is comparable with the value we obtained after 2-3 s of shading, and also comparable with our shaded values for *pgm* and *adg1*. Thus, it seems possible that the similar values for wild-type and mutant plants reported by Bahaji et al. (2014) do not reflect the true levels of ADPG in illuminated leaves, explaining the marked difference from our results showing that WT plants have much higher levels of ADPG than the *pgm* and *adg1* mutants when leaves are sampled in a way that preserves the *in-vivo* levels in the light.

Our metabolite data from WT and mutant (*pgm* and *adg1*) plants conflict with equivalent data that have been used to challenge the consensus pathway of starch synthesis. Our data are entirely consistent with predictions based on the consensus pathway being the predominant, if not the only, pathway of starch synthesis in leaves. Given the precautions we took to ensure that *in-vivo* metabolite levels were preserved during tissue sampling and the robustness of the LC-MS/MS assay for ADPG, we would argue that our results are the more trustworthy.

To test whether the proposed alternative pathway of ADPG synthesis, via SUS, makes any significant contribution to starch synthesis, we also re-analyzed the *sus1234* mutant reported in Barratt et al. (2009) and generated sextuple *sus123456* mutants that lack all known isoforms of SUS in Arabidopsis. The *sus1234* mutant lacks *SUS* expression in mesophyll cells where transitory starch is made in WT plants (Barratt et al., 2009; Smith et al., 2012) and lacks soluble SUS activity in siliques, where activity is high and readily measurable in WT plants(Fig. 4A). The *sus1234* mutant should have less starch than WT plants if the SUS-pathway supplies a significant amount of ADPG for starch synthesis. However, we confirmed that starch levels in this mutant are indistinguishable from WT, as reported in Barratt et al. (2009) and Baroja-Fernández et al. (2012a), and showed that the mutant also has WT levels of ADPG and of the two other intermediates of the consensus pathway, Glc6P and Glc1P (Figs 3 and 5). Likewise, two sextuple *sus123456* mutants that had no detectable SUS activity contained essentially WT levels of key intermediates from the consensus pathways for synthesis of starch (Glc6P, Glc1P, ADPG) and sucrose (DHAP, Fru6P, UDPG), as well as WT levels of starch and soluble sugars (Fig. 5, Supplemental Fig. S4). These results argue strongly against the proposed SUS pathway making any significant contribution to ADPG and starch synthesis in leaves. The sextuple *sus123456* mutants showed near WT-like growth, indicating that sucrose synthase is not an essential enzyme in Arabidopsis, at least under standard laboratory growth conditions. However, SUS may contribute to fitness in natural environments, especially if the plants are subjected to flooding or hypoxic stress (Bieniewska et al., 2007).

## MATERIALS AND METHODS

### Plant materials

Arabidopsis (*Arabidopsis thaliana* [L.] Heynh) Columbia-0 wild-type and the *pgm* (Caspar et al., 1985), *adg1* (Lin et al., 1988a) and *sus1234* (Barratt et al., 2009) mutant germplasm were from in-house collections.

Sextuple *sus123456* mutants were generated by gene editing of the *SUS5* and *SUS6* genes in the *sus1234* (Barratt et al., 2009) background using CRISPR/Cas9 and two pairs of guide RNA (gRNA) targets: *SUS5* – GAAATGACATCTGGATCGTT and TGTAGAACTTGGTGAATCTC; *SUS6* – GGTAAGGGTAGATATCGAAT and GTTCTTGAAGCACCAGACAA. The CRISPR/Cas9 and gRNA sequences were cloned into the pHEE401E vector as described in Xin et al. (2014) and Wang et al. (2015). The construct was introduced into the *sus1234* quadruple mutant by agrobacterium-mediated transformation using the floral-dip method (Clough and Bent, 1998) to generate sextuple *sus123456* mutants. The CRISPR/Cas9 construct was subsequently eliminated by crossing the sextuple *sus123456* lines with the quadruple *sus1234* parent. Progeny were screened by PCR to identify lines that were homozygous for the T-DNA insertions in *SUS1-SUS4* and for gene-edited mutant alleles of *SUS5* and *SUS6* (Supplemental Fig. S2). Mutations in the *SUS5* gene were identified by PCR using primers sus5-Fw and sus5-Rv (Supplemental Table S3) and restriction with HinfI. Mutations in the *SUS6* gene were identified by PCR using primers sus6-Fw and sus6-Rv (Supplemental Table S3) and restriction with Van91I (Supplemental Fig. S2).

Plants were grown in a 1:1 mixture of compost and vermiculite in 6-cm diameter pots with a 16-h photoperiod (140 μE m^−2^ s^−1^ irradiance provided by white fluorescent tubes) and day/night temperatures of 20°C/18°C.

### Enzyme activity measurements

Finely ground frozen leaf tissue (20 mg) was homogenized in 1 mL of ice-cold extraction buffer as described in Gibon et al. (2004). After centrifugation of the crude extract at 10,000 x *g* (4°C) for 5 min, an aliquot (180 μL) of the supernatant was desalted by centrifugation through a MicroSpin column (GE healthcare Life sciences, Marlborough, MA, USA) equilibrated with extraction buffer. Soluble SUS activity was determined in the sucrose cleavage direction by measuring the UDP-dependent production of UDPG from sucrose, with enzymatic determination of UDPG as described by Dancer and ap Rees (1989). Reaction mixtures (225 μl) contained: 20 mM PIPES-KOH pH 6.5, 3 mM MgCl_2_, 100 mM sucrose and 2 mM UDP, with UDP being omitted from blank reactions. Reactions were started by addition of 25 μl of desalted extract and incubation at 30°C for 25 min. The reaction was stopped by addition of 250 μL of 50 mM Tricine-KOH, pH 8.3, and heating at 100°C for 2 min. UDPG was measured enzymatically by coupling to reduction of NADP and monitoring the increase in absorbance at 340 nm. The initial reaction mixture (500 μL) contained: 50 mM Tricine-HCl, pH 8.0, 4 mM MgCl_2_, 12.5 mM NADP, 0.3 unit glucose-6-phosphate dehydrogenase (EC 1.1.1.49; from yeast), 0.4 unit phosphoglucomutase (EC 5.4.2.2; from rabbit muscle) and 400 μl of the SUS assay reaction. When the A_340_ was stable, following consumption of glucose 6-phosphate and glucose 1-phosphate, UDPG was determined by addition of sodium pyrophosphate (final concentration 1 mM) and 0.2 U UDP-glucose pyrophosphorylase (EC 2.7.7.9; from yeast). All coupling enzymes were from Sigma-Aldrich (https://www.sigmaaldrich.com). ASPP activity was measured as described in Baroja-Fernández et al. (2004), using ADP-ribose or ADPG as substrates, except that the products of the reaction (ribose 5-phosphate and Glc1P, respectively) were measured by LC-MS/MS (Arrivault et al., 2009).

### Metabolite measurements

Rosettes were rapidly quenched in the light by either pouring liquid nitrogen directly onto the rosettes, or by cutting the hypocotyl from below and rapidly transferring the cut rosette into liquid nitrogen in the light, taking care not to shade the leaves at any time. Phosphorylated intermediates were extracted using chloroform-methanol as described in Lunn et al. (2006) and measured by ion-pair reverse phase LC-MS/MS (Arrivault et al., 2009) or anion-exchange LC-MS/MS (Lunn et al., 2006; with modifications as described in Figueroa et al., 2016). Soluble sugars were measured in ethanolic extracts and starch was measured in the ethanol-insoluble residue as described in (Stitt et al., 1989).

## ACKNOWLEDGEMENTS

We thank Christin Abel for help with plant cultivation. This work was supported by a PhD fellowship (to M.M.F.F.F.) from the International Max Planck Research School “Primary Metabolism and Plant Growth), the European Commission FP7 collaborative project TiMet (contract No. 245142, to A.M.S. and M.S.), the Swedish Research Council for Sustainable Development (Formas) (project grant 2016-01322 to T.N.) and the Max Planck Society (H.I., S.A., R.F., M.S., and J.E.L.).

## AUTHOR CONTRIBUTIONS

W.W. generated the sextuple *sus* mutants and performed the morphological phenotyping. M.M.F.F.F. performed the harvesting trials and, with H.I., collected samples for metabolite analysis. M.M.F.F.F., S.A., H.I. and R.F. measured metabolites. H.I. measured SUS activity. T.N., M.S., A.M.S. and J.E.L. conceived the study. J.E.L. and T.N. drafted the manuscript.

---

**Supplemental Figure S1.** Schematic diagram of the gene edited mutations in the Arabidopsis *SUS5* and *SUS6* genes.

**Supplemental Figure S2.** PCR genotyping of Arabidopsis *sus* mutants.

**Supplemental Figure S3.** Expression atlas of Arabidopsis *SUS1-SUS6* transcripts.

**Supplemental Figure S4.** Metabolite levels in wild-type Arabidopsis and *sus* mutants.

**Supplemental Table S1.** Statistical analysis of metabolite data in Figure 3.

**Supplemental Table S2.** Statistical analysis of metabolite data in Figure 5 and Supplemental Figure S4.

**Supplemental Table S3.** Oligonucleotide primers for PCR genotyping.

## Notes

### Competing Interest Statement

The authors have declared no competing interest.

## REFERENCES

Arrivault S, Guenther M, Florian A, Encke B, Feil R, Vosloh D, Lunn JE, Sulpice R, Fernie AR, Stitt M, Schulze WX (2014) Dissecting the subcellular compartmentation of proteins and metabolites in Arabidopsis leaves using non-aqueous fractionation. Mol Cell Proteom 13: 2246–2259.

Arrivault S, Guenther M, Ivakov A, Feil R, Vosloh D, van Dongen JT, Sulpice R, Stitt M (2009) Use of reverse-phase liquid chromatography, linked to tandem mass spectrometry, to profile the Calvin cycle and other metabolic intermediates in Arabidopsis rosettes at different carbon dioxide concentrations. Plant J 59: 826–839.

Bahaji A, Baroja-Fernández E, Sánchez-López AM, Muñoz FJ, Li J, Almagro G, Montero M, Pujol P, Galarza R, Kaneko K, Oikawa K, Wada K, Mitsui T, Pozueta-Romero J (2014) HPLC-MS/MS analyses show that the near-starchless *aps1* and *pgm* leaves accumulate wild type levels of ADPglucose: further evidence for the occurrence of important ADPglucose biosynthetic pathway(s) alternative to the pPGI-pPGM-AGP pathway. PLoS One 9: e104997.

Bahaji A, Li J, Ovecka M, Ezquer I, Muñoz FJ, Baroja-Fernández E, Romero JM, Almagro G, Montero M, Hidalgo M, Sesma MT, Pozueta-Romero J (2011) *Arabidopsis thaliana* mutants lacking ADP-glucose pyrophosphorylase accumulate starch and wild-type ADP-glucose content: further evidence for the occurrence of important sources, other than ADP-glucose pyrophosphorylase, of ADP-glucose linked to leaf starch biosynthesis. Plant Cell Physiol 52: 1162–1176.

Bahaji A, Sánchez-López ÁM, De Diego N, Muñoz FJ, Baroja-Fernández E, Li J, Ricarte-Bermejo A, Baslam M, Aranjuelo I, Almagro G, Humplík JF, Novák O, Spíchal L, Doležal K, Pozueta-Romero J (2015) Plastidic phosphoglucose isomerase is an important determinant of starch accumulation in mesophyll cells, growth, photosynthetic capacity, and biosynthesis of plastidic cytokinins in Arabidopsis. PLoS One 10: e0119641.

Bahaji A, Almagro G, Ezquer I, Gámez-Arcas S, Sánchez-López ÁM, Muñoz FJ, Barrio RJ, Sampedro MC, De Diego N, Spíchal L, Doležal K, Tarkowská D, Caporali E, Mendes MA, Baroja-Fernández E, Pozueta-Romero J (2018) Plastidial phosphoglucose isomerase is an important determinant of seed yield through its involvement in gibberellin-mediated reproductive development and storage reserve biosynthesis in Arabidopsis. Plant Cell 30: 2082–2098.

Ballicora MA, Iglesias AA, Preiss J (2004) ADP-glucose pyrophosphorylase: a regulatory enzyme for plant starch synthesis. Photosynth Res 79:1–24.

Baroja-Fernández E, Munoz FJ, Akazawa T, Pozueta-Romero J (2001) Reappraisal of the currently prevailing model of starch biosynthesis in photosynthetic tissues: a proposal involving the cytosolic production of ADP-glucose by sucrose synthase and occurrence of cyclic turnover of starch in the chloroplast. Plant Cell Physiol 42: 1311–1320.

Baroja-Fernández E, Muñoz F J, Saikusa T, Rodríguez-López M, Akazawa T, Pozueta-Romero J (2003) Sucrose synthase catalyzes the de novo production of ADPglucose linked to starch biosynthesis in heterotrophic tissues of plants. Plant Cell Physiol 44: 500–509.

Baroja-Fernández E, Muñoz F J, Zandueta-Criado A, Morán-Zorzano MT, Viale AM, Alonso-Casajús N, Pozueta-Romero J (2004) Most of ADP-glucose linked to starch biosynthesis occurs outside the chloroplast in source leaves. Proc Nat Acad Sci USA 101: 13080–13085.

Baroja-Fernández E, Muñoz FJ, Pozueta-Romero J (2005) Response to Neuhaus et al.: No need to shift the paradigm on the metabolic pathway to transitory starch in leaves. Trends Plant Sci 10: 156–158.

Baroja-Fernández E, Muñoz FJ, Li J, Bahaji A, Almagro G, Montero M, Etxeberria E, Hidalgo M, Sesma MT, Pozueta-Romero J (2012a) Sucrose synthase activity in the *sus1/sus2/sus3/sus4 Arabidopsis* mutant is sufficient to support normal cellulose and starch production. Proc Nat Acad Sci USA 109: 321–326.

Baroja-Fernández E, Muñoz FJ, Bahaji A, Almagro G, Pozueta-Romero J (2012b) Reply to Smith et al.: No evidence to challenge the current paradigm on starch and cellulose biosynthesis involving sucrose synthase activity. Proc Nat Acad Sci USA 109: E777.

Barratt DHP, Derbyshire P, Findlay K, Pike M, Wellner N, Lunn J, Feil R, Simpson C, Maule A J, Smith AM (2009) Normal growth of Arabidopsis requires cytosolic invertase but not sucrose synthase. Proc Nat Acad Sci USA 106: 13124–13129.

Beckles DM, Smith AM, ap Rees T (2001) A cytosolic ADP-glucose pyrophosphorylase is a feature of graminaceous endosperms, but not of other starch-storing organs. Plant Physiol 125: 818–827.

Bieniawska Z, Paul Barratt DH, Garlick AP, Thole V, Kruger NJ, Martin C, Zrenner R, Smith AM (2007) Analysis of the sucrose synthase gene family in Arabidopsis. Plant J 49: 810–828.

Bowsher CG, Scrase-Field EFAL, Esposito S, Emes MJ, Tetlow IJ (2007) Characterization of ADP-glucose transport across the cereal endosperm amyloplast envelope. J Exp Bot 58: 1321–1332.

Caspar T, Huber SC, Somerville C (1985) Alterations in growth, photosynthesis, and respiration in a starchless mutant of *Arabidopsis thaliana* (L.) deficient in chloroplast phosphoglucomutase activity. Plant Physiol 79: 11–17.

Clough SJ, Bent AF (1998) Floral dip: a simplified method for Agrobacterium-mediated transformation of *Arabidopsis thaliana*. Plant J 16: 735–743.

Colleoni C, Linka M, Deschamps P, Handford MG, Dupree P, Weber AP, Ball SG (2010) Phylogenetic and biochemical evidence supports the recruitment of an ADPglucose translocator for the export of photosynthate during plastid endosymbiosis. Mol Biol Evol 27: 2691–2701.

Dancer JE, ap Rees T (1989) Relationship between pyrophosphate: fructose-6-phosphate 1-phosphotransferase, sucrose breakdown, and respiration. J Plant Physiol 135:197–206.

Dawson RMC, Elliott DC, Elliott WH, Jones KM (1986) Data for Biochemical Research, third edition (Oxford University Press, Oxford), pp103–114.

Denyer K, Dunlap F, Thorbjørnsen T, Keeling P, Smith AM (1996) The major form of ADP-glucose pyrophosphorylase in maize endosperm is extra-plastidial. Plant Physiol 112: 779–785.

Emes MJ, Bowsher CG, Hedley C, Burrell MM, Scrase-Field ESF, Tetlow IJ (2003) Starch synthesis and carbon partitioning in developing endosperm. J Exp Bot 54: 569–575.

Fallahi H, Scofield GN, Badger MR, Chow WS, Furbank RT, Ruan YL (2008) Localization of sucrose synthase in developing seed and siliques of *Arabidopsis thaliana* reveals diverse roles for SUS during development. J Exp Bot 59: 3283–3295.

Fernandez O, Ishihara H, George GM, Mengin V, Flis A, Sumner D, Arrivault S, Feil R, Lunn JE, Zeeman SC, Smith AM, Stitt M (2017) Leaf starch turnover occurs in long days and in falling light at the end of the day. Plant Physiol 174: 2199–2212.

Fettke J, Malinova I, Albrecht T, Hejazi M, Steup M (2011) Glucose-1-phosphate transport into protoplasts and chloroplasts from leaves of Arabidopsis. Plant Physiol 155: 1723–1734.

Figueroa CM, Feil R, Ishihara H, Watanabe M, Kölling K, Krause U, Höhne M, Encke B, Plaxton WC, Zeeman SC, Li Z, Schulze WX, Hoefgen R, Stitt M, Lunn JE (2016) Trehalose 6-phosphate coordinates organic and amino acid metabolism with carbon availability. Plant J 85: 410–423.

Gibon, Y., Blaesing, O.E., Hannemann, J., Carillo, P., Höhne, M., Hendriks, J.H.M., Palacios, N., Cross, J., Selbig, J., Stitt, M. (2004). A robot-based platform to measure multiple enzyme activities in Arabidopsis using a set of cycling assays: comparison of changes of enzyme activities and transcript levels during diurnal cycles and in prolonged darkness. Plant Cell 16, 3304–3325.

Hädrich N, Hendriks JH, Kötting O, Arrivault S, Feil R, Zeeman SC, Gibon Y, Schulze WX, Stitt M, Lunn JE (2012) Mutagenesis of cysteine 81 prevents dimerization of the APS1 subunit of ADP-glucose pyrophosphorylase and alters diurnal starch turnover in *Arabidopsis thaliana* leaves. Plant J 70: 231–242.

Hädrich N, Gibon Y, Schudoma C, Altmann T, Lunn JE, Stitt M (2011) Use of TILLING and robotised enzyme assays to generate an allelic series of *Arabidopsis thaliana* mutants with altered ADP-glucose pyrophosphorylase activity. J Plant Physiol 168: 1395–1405.

Heldt HW, Chon CJ, Maronde D, Herold A, Stankovic ZS, Walker DA, Kraminer A, Kirk MR, Heber U (1977) Role of orthophosphate and other factors in the regulation of starch formation in leaves and isolated chloroplasts. Plant Physiol 59: 1146–1155.

Hill BL, Figueroa CM, Ascencion MD, Lunn JE, Iglesias AA, Ballicora MA (2017) On the stability of nucleoside diphosphate glucose metabolites: implications for studies of plant carbohydrate metabolism. J Exp Bot 68: 3331–3337.

Ishihara H, Obata T, Sulpice R, Fernie AR, Stitt M (2015) Quantifying protein synthesis and degradation in Arabidopsis by dynamic ^13^CO2 labeling and analysis of enrichment in individual amino acids in their free pools and in protein. Plant Physiol. 168: 74–93.

Kirchberger S, Tjaden J, Neuhaus HE (2008) Characterization of the Arabidopsis Brittle1 transport protein and impact of reduced activity on plant metabolism. Plant J 56: 51–63.

Lin TP, Caspar T, Somerville C, Preiss J (1988a) Isolation and characterization of a starchless mutant of *Arabidopsis thaliana* (L.) Heynh lacking ADPglucose pyrophosphorylase activity. Plant Physiol 86: 1131–1135.

Lin TP, Caspar T, Somerville CR, Preiss J (1988b) A starch deficient mutant of *Arabidopsis thaliana* with low ADPglucose pyrophosphorylase activity lacks one of the two subunits of the enzyme. Plant Physiol 88: 1175–1181.

Lunn JE, Feil R, Hendriks JH, Gibon Y, Morcuende R, Osuna D, Scheible WR, Carillo P, Hajirezaei MR, Stitt M (2006) Sugar-induced increases in trehalose 6-phosphate are correlated with redox activation of ADPglucose pyrophosphorylase and higher rates of starch synthesis in *Arabidopsis thaliana*. Biochem J 397: 139–148.

Malinova I, Kunz HH, Alseekh S, Herbst K, Fernie AR, Gierth M, Fettke J (2014) Reduction of the cytosolic phosphoglucomutase in Arabidopsis reveals impact on plant growth, seed and root development, and carbohydrate partitioning. PLoS One 9: e112468.

Moreno-Bruna B, Baroja-Fernández E, Muñoz FJ, Bastarrica-Berasategui A, Zandueta-Criado A, Rodríguez-López M, Lasa I, Akazawa T, Pozueta-Romero J (2001) Adenosine diphosphate sugar pyrophosphatase prevents glycogen biosynthesis in *Escherichia coli*. Proc Nat Acad Sci USA 98: 8128–8132.

Muñoz FJ, Baroja-Fernández E, Morán-Zorzano MT, Alonso-Casajús N, Pozueta-Romero J (2006) Cloning, expression and charcterization of a nudix hydrolase that catalyzes the hydrolytic breakdown of ADP-glucose linked to starch biosynthesis in *Arabidopsis thaliana*. Plant Cell Physiol 47: 926–934.

Muñoz FJ, Baroja-Fernández E, Morán-Zorzano MT, Viale AM, Etxeberria E, Alonso-Casajús N, Pozueta-Romero J (2005) Sucrose synthase controls both intracellular ADP glucose levels and transitory starch biosynthesis in source leaves. Plant Cell Physiol 46: 1366–1376.

Neuhaus HE, Häusler RE, Sonnewald U (2005) No need to shift the paradigm on the metabolic pathway to transitory starch in leaves. Trends Plant Sci 10: 154–156.

Neuhaus HE, Stitt M (1990) Control analysis of photosynthate partitioning – impact of reduced activity of ADP-glucose pyrophosphorylase or plastid phosphoglucomutase on the fluxes to starch and sucrose in *Arabidopsis thaliana* (L.) Heynh. Planta 182: 445–454.

Neuhaus HE, Thom E, Möhlmann T, Steup M, Kampfenkel K (1997) Characterization of a novel eukaryotic ATP/ADP translocator located in the plastid envelope of *Arabidopsis thaliana* L. Plant J 11: 73–82.

Niittylä T, Messerli G, Trevisan M, Chen J, Smith AM, Zeeman SC (2004) A previously unknown maltose transporter essential for starch degradation in leaves. Science 303: 87–89.

Okita TW (1992) Is there an alternative pathway for starch synthesis? Plant Physiol 100: 560–564.

Okita TW, Greenberg E, Kuhn DN, Preiss J (1979) Subcellular localization of the starch degradative and biosynthetic enzymes of spinach leaf. Plant Physiol 64: 187–192.

Perata P, Pozueta-Romero J, Yamaguchi J, Akazawa T (1992) Artifactual detection of ADP-dependent sucrose synthase in crude plant extracts. FEBS Letts 309: 283–297.

Pozueta-Romero J, Yamaguchi J, Akazawa T (1991) ADPG formation by the ADP-specific cleavage of sucrose – reassessment of sucrose synthase. FEBS Letts 291: 233–237.

Rodriguez-López M, Baroja-Fernàndez E, Zandueta-Criado A, Pozueta-Romero J (2000) Adenosine diphosphate glucose pyrophosphatase: a plastidial phosphodiesterase that prevents starch biosynthesis. Proc Nat Acad Sci USA 97: 8705–8710.

Schneider A, Häusler RE, Kolukisaoglu U, Kunze R, van der Graaff E, Schwacke R, Catoni E, Desimone M, Flügge U-I (2002) An *Arabidopsis thaliana* knock-out mutant of the chloroplast triose phosphate/phosphate translocator is severely compromised only when starch synthesis, but not starch mobilisation is abolished. Plant J 32: 685–699.

Shannon JC, Pien FM, Cao H, Liu KC (1998) Brittle-1, an adenylate translocator, facilitates transfer of extraplastidial synthesized ADP-glucose into amyloplasts of maize endosperms. Plant Physiol 117:1235–1252.

Strand A, Zrenner R, Trevanion S, Stitt M, Gustafsson P, Gardeström P (2000) Decreased expression of two key enzymes in the sucrose biosynthesis pathway, cytosolic fructose-1,6-bisphosphatase and sucrose phosphate synthase, has remarkably different consequences for photosynthetic carbon metabolism in transgenic *Arabidopsis thaliana*. Plant J 23: 759–770.

Sun J, Zhang J, Larue CT, Huber SC (2011) Decrease in leaf sucrose synthesis leads to increased leaf starch turnover and decreased RuBP regeneration-limited photosynthesis but not Rubisco-limited photosynthesis in Arabidopsis null mutants of SPSA1. Plant Cell Environ 34: 592–604.

Szecowka M, Heise R, Tohge T, Nunes-Nesi A, Vosloh D, Huege J, Feil R, Lunn J, Nikoloski Z, Stitt M, Fernie AR, Arrivault S (2013) Metabolic fluxes in an illuminated Arabidopsis rosette. Plant Cell 25: 694–714.

Smith AM, Kruger NJ, Lunn JE (2012) Source of sugar nucleotides for starch and cellulose synthesis. Proc Nat Acad Sci USA 109: E776.

Smith AM, Zeeman SC (2020) Starch: a flexible, adaptable carbon store coupled to plant growth. Ann Rev Plant Biol 71: 1–29.

Stitt M, Lilley RM, Gerhardt R, Heldt H-W (1989) Metabolite levels in specific cells and subcellular compartments of plant-leaves. Meth Enzymol 174: 518–552.

Stitt M, Zeeman SC (2012) Starch turnover: pathways, regulation and role in growth. Curr Opin Plant Biol 15: 282–292.

Streb S, Egli B, Eicke S, Zeeman SC (2009) The debate on the pathway of starch synthesis: A closer look at low-starch mutants lacking plastidial phosphoglucomutase supports the chloroplast-localized pathway. Plant Physiol 151: 1769–1772.

Sun J, Okita TW, Edwards GE (1999) Modification of carbon partitioning, photosynthetic capacity, and O2 sensitivity in Arabidopsis plants with low ADP-glucose pyrophosphorylase activity. Plant Physiol 119: 267–276.

Thorbjørnsen T, Villand P, Denyer K, Olsen OA, Smith AM (1996) Distinct isoforms of ADPglucose pyrophosphorylase occur inside and outside the amyloplasts in barley endosperm. Plant J 10: 243–250.

Wang ZP, Xing HL, Dong L, Zhang HY, Han CY, Wang XC, Chen QJ (2015) Egg cell-specific promoter-controlled CRISPR/Cas9 efficiently generates homozygous mutants for multiple target genes in Arabidopsis in a single generation. Genome Biol 2015;16:144.

Xing HL, Dong L, Wang ZP, Zhang HY, Han CY, Liu B, Wang XC, Chen QJ (2014) A CRISPR/Cas9 toolkit for multiplex genome editing in plants. BMC Plant Biol 14: 327.

Yao D, Gonzales-Vigil E, Mansfield SD (2020) Arabidopsis sucrose synthase localization indicates a primary role in sucrose translocation in phloem. J Exp Bot 71: 1858–1869.

Yu TS, Lue WL, Wang SM, Chen J (2000) Mutation of Arabidopsis plastid phosphoglucose isomerase affects leaf starch synthesis and floral initiation. Plant Physiol 123: 319–326.

