## Supplemental Data for "The pathway of starch synthesis in *Arabidopsis thaliana* leaves"

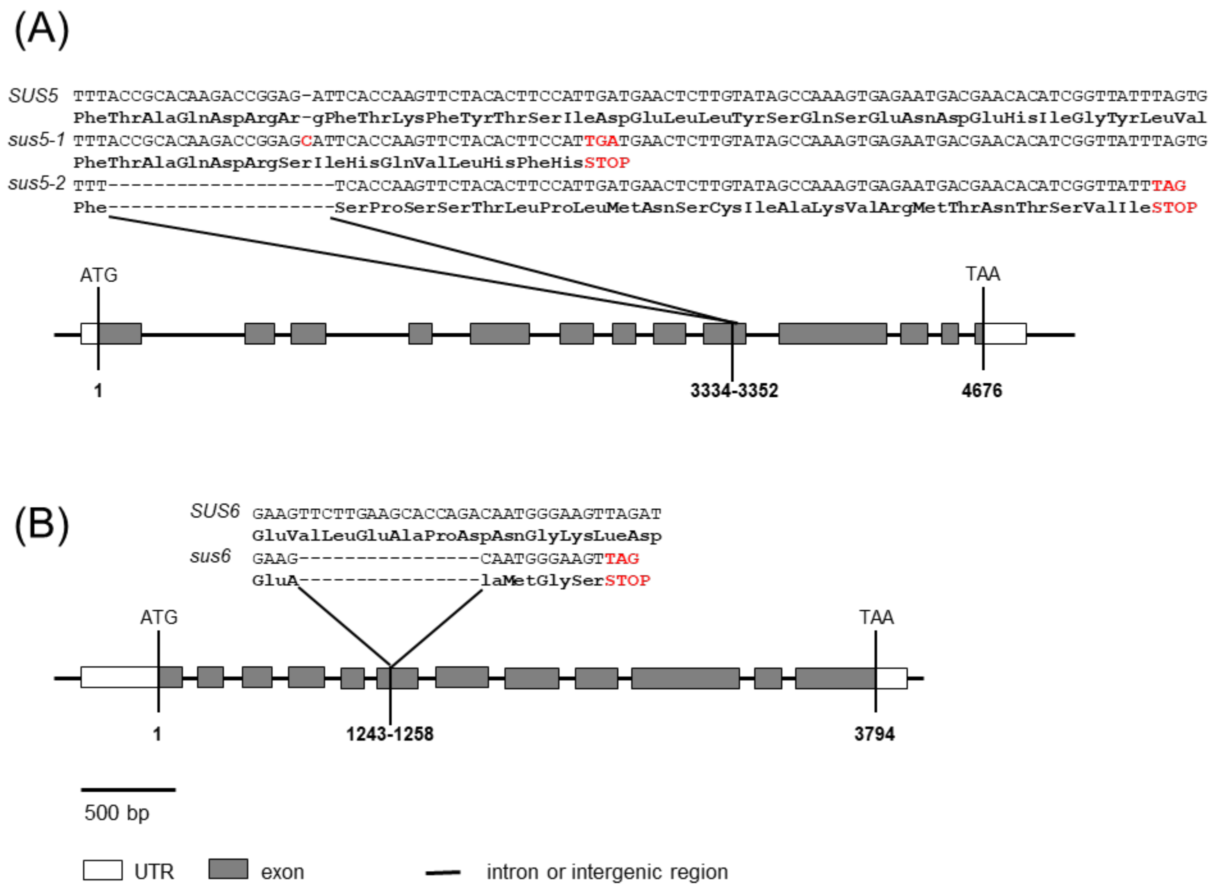


**Supplemental Figure S1. Schematic diagram of the gene edited mutations in the Arabidopsis *SUS5* and *SUS6* genes.**

(A) *SUS5* gene showing the wild-type (Col-0) sequence, the *sus5-1* allele containing a single base (C) insertion (shown in red) and the *sus5-2* allele containing a 20-bp deletion. (B) *SUS6* gene showing the wild-type (Col-0) sequence and the *sus6* allele containing a 16-bp deletion. All mutations generated premature STOP codons (shown in red). The dark grey and white boxes indicate translated and untranslated regions, respectively.

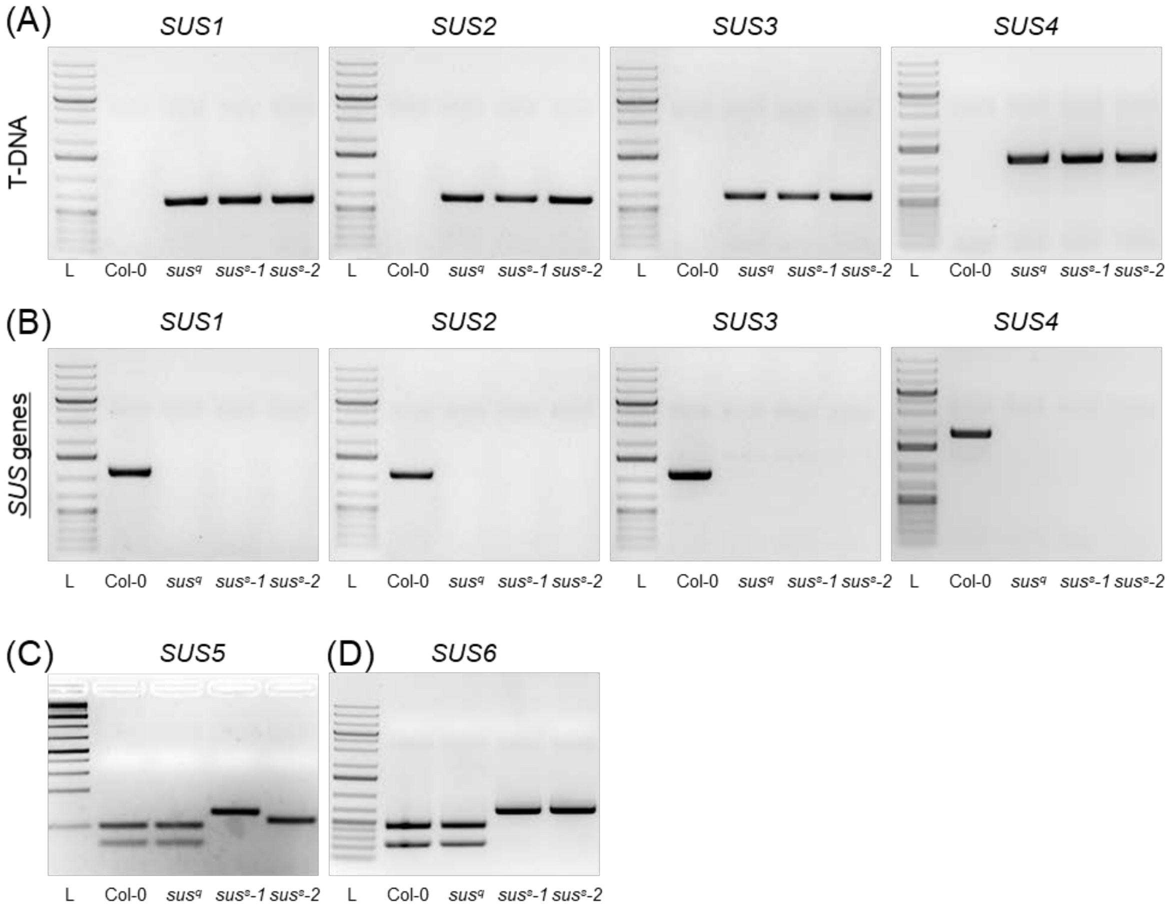


**Supplemental Figure S2. PCR genotyping of Arabidopsis *sus* mutants.**

(A) T-DNA insertions in the *SUS1*, *SUS2*, *SUS3*and *SUS4* genes were confirmed by PCR using gene specific (LP) and T-DNA border primers (LBb1.3 for *sus1*, *sus2*, *sus3* and P66 for *sus4*). (B) Homozygosity of T-DNA insertion loci in the *SUS1*, *SUS2*, *SUS3* and *SUS4* genes was confirmed by genomic PCR using *SUS* gene specific primers (LP and RP) that flank the respective T-DNA insertions. (C) Identification of mutations in the *SUS5* gene by genomic PCR (using primers sus5-Fw and sus5-Rv) and restriction with HinfI. (D) Identification of mutations in the *SUS6* gene by genomic PCR (using primers sus6-Fw and sus6-Rv) and restriction with Van91I. Abbreviations: L, 1-kb plus ladder (Thermo Scientific); Col-0, wild type Columba-0; *Sus*$^{q}$, *sus1234* quadruple mutant; *sus*$^{s}$*-1*, *sus12345*$^{1}$*6* sextuple mutant; *sus*$^{s}$*-2*, *sus12345*$^{2}$*6* sextuple mutant.

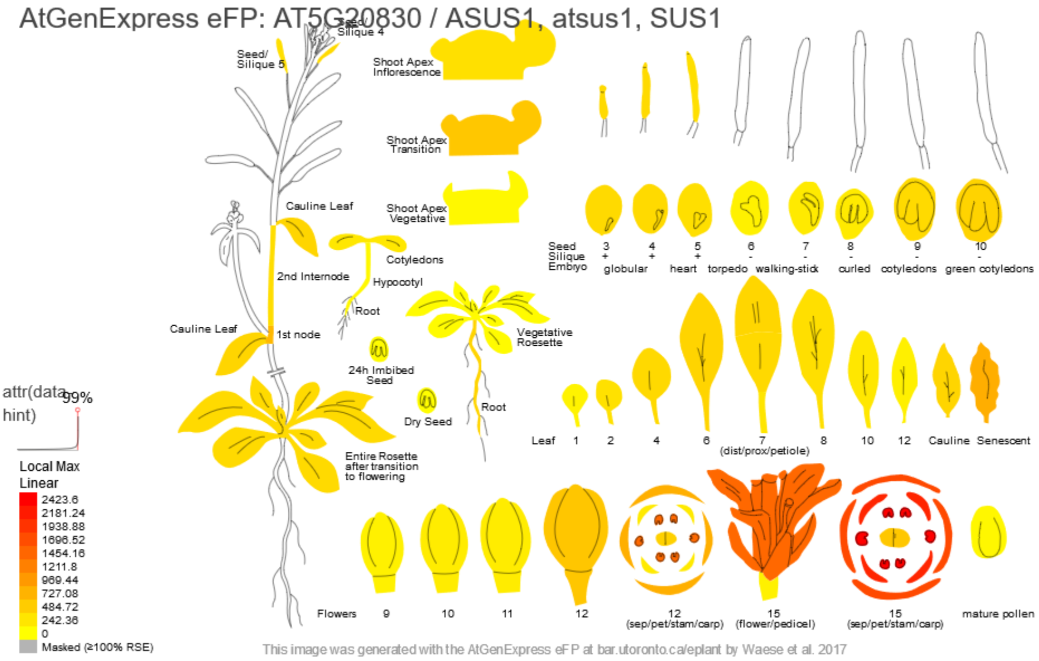


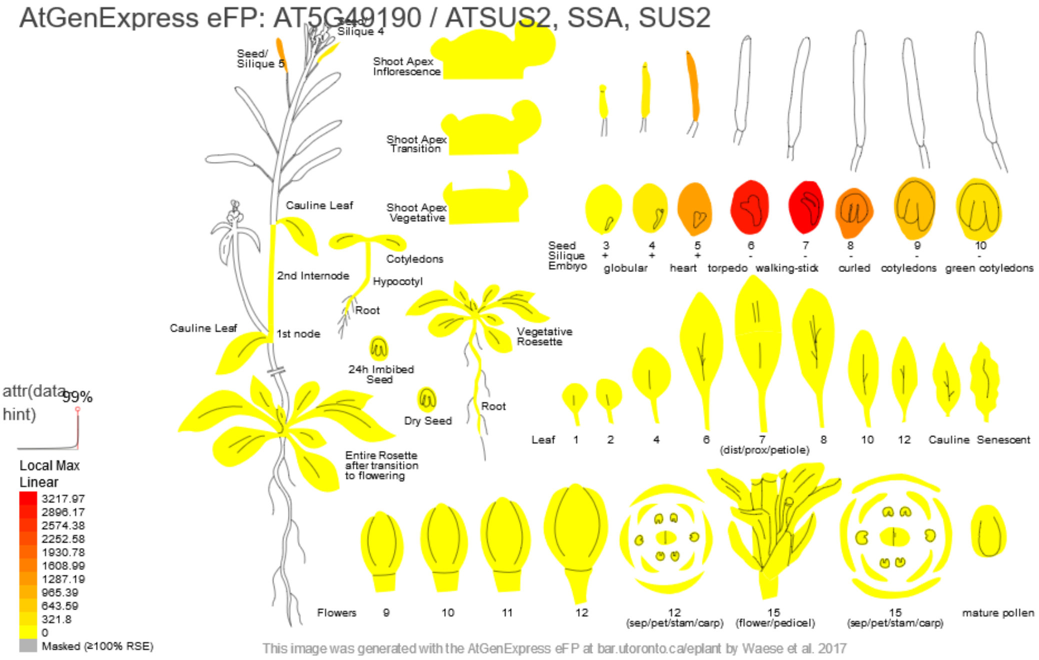


**Supplemental Figure S3a. Expression atlas of Arabidopsis *SUS1* and *SUS2* transcripts.**

*SUS1* (At5g20830) and *SUS2* (At5g49190) transcript abundance in different Arabidopsis tissues and developmental stages based on Affymetrix ATH1 array data from Schmid et al. (2005), and visualized with the Plant eFP browser (bar.utoronto.ca/eplant; Waese et al., 2016).

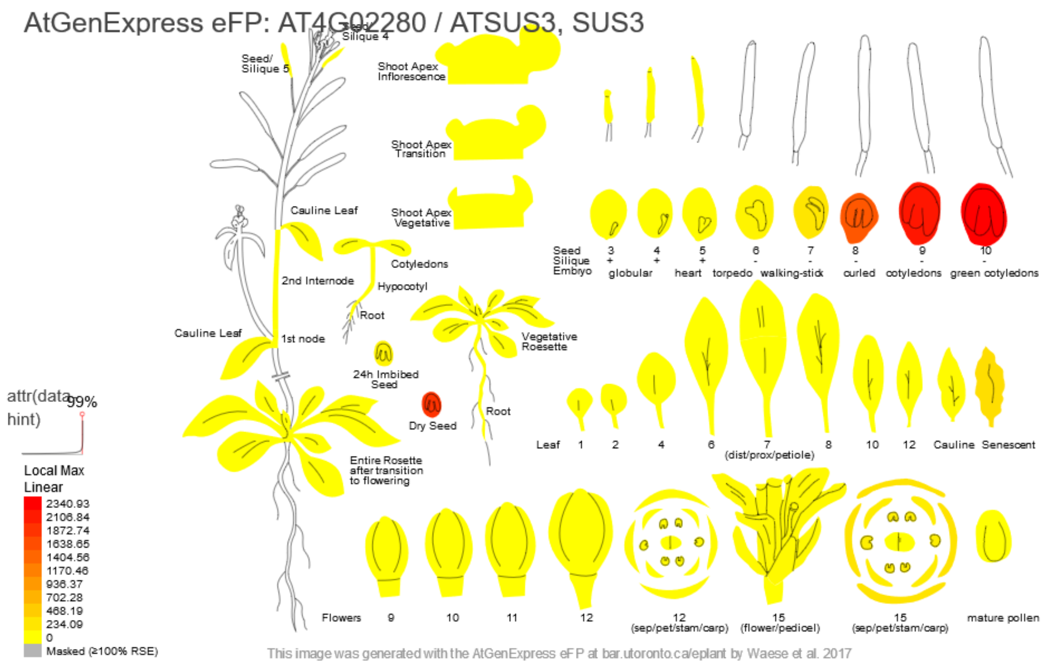


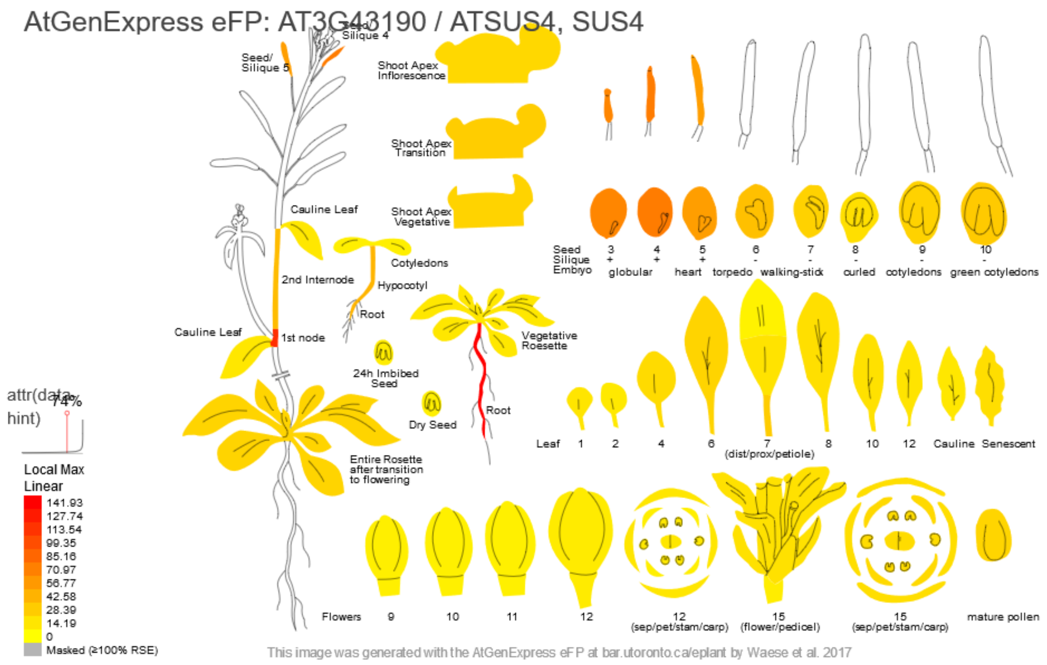


**Supplemental Figure S3b. Expression atlas of Arabidopsis *SUS3* and *SUS4* transcripts.**

*SUS3* (At4g02280) and *SUS4* (At3g43190) transcript abundance in different Arabidopsis tissues and developmental stages based on Affymetrix ATH1 array data from Schmid et al. (2005), and visualized with the Plant eFP browser (bar.utoronto.ca/eplant; Waese et al., 2016).

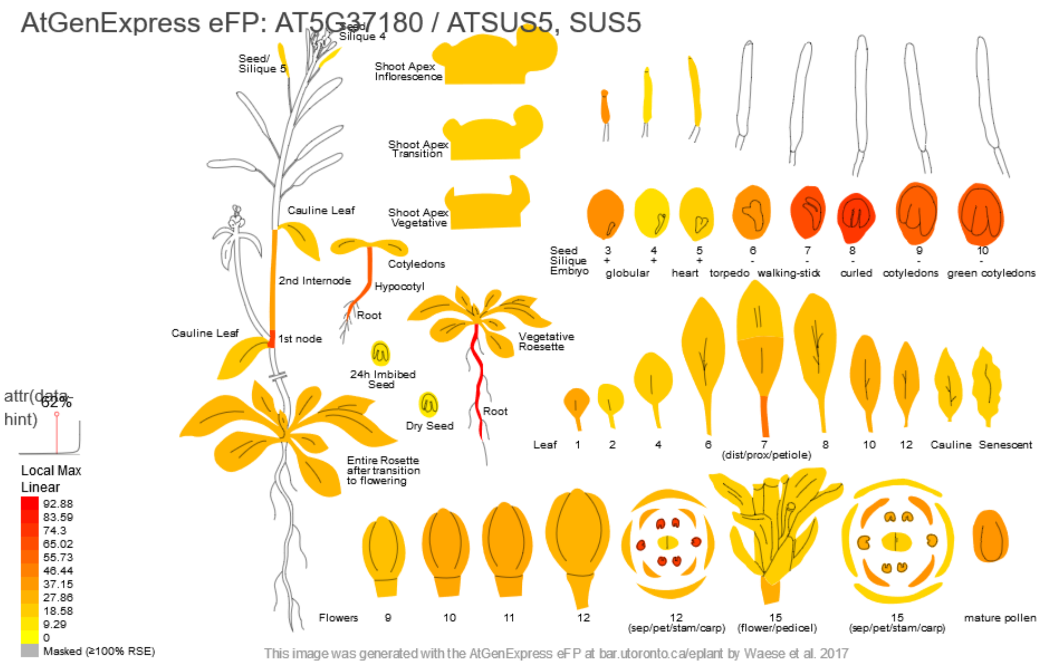


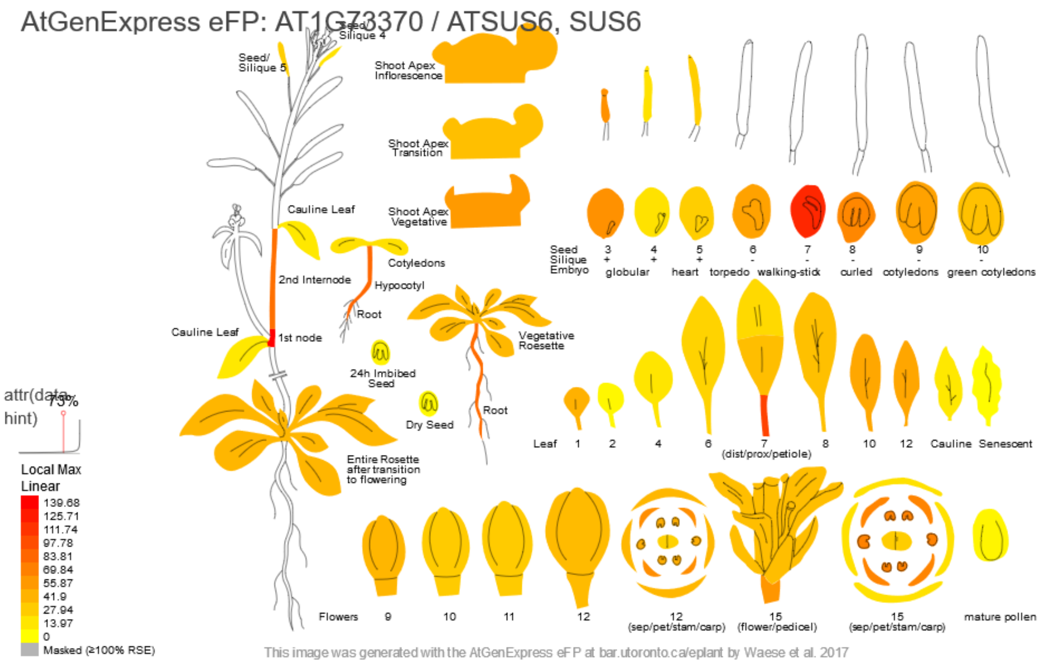


**Supplemental Figure S3c. Expression atlas of Arabidopsis *SUS5* and *SUS6* transcripts.**

*SUS5* (At5g37180) and *SUS6* (At1g73370) transcript abundance in different Arabidopsis tissues and developmental stages based on Affymetrix ATH1 array data from Schmid et al. (2005), and visualized with the Plant eFP browser (bar.utoronto.ca/eplant; Waese et al., 2016).

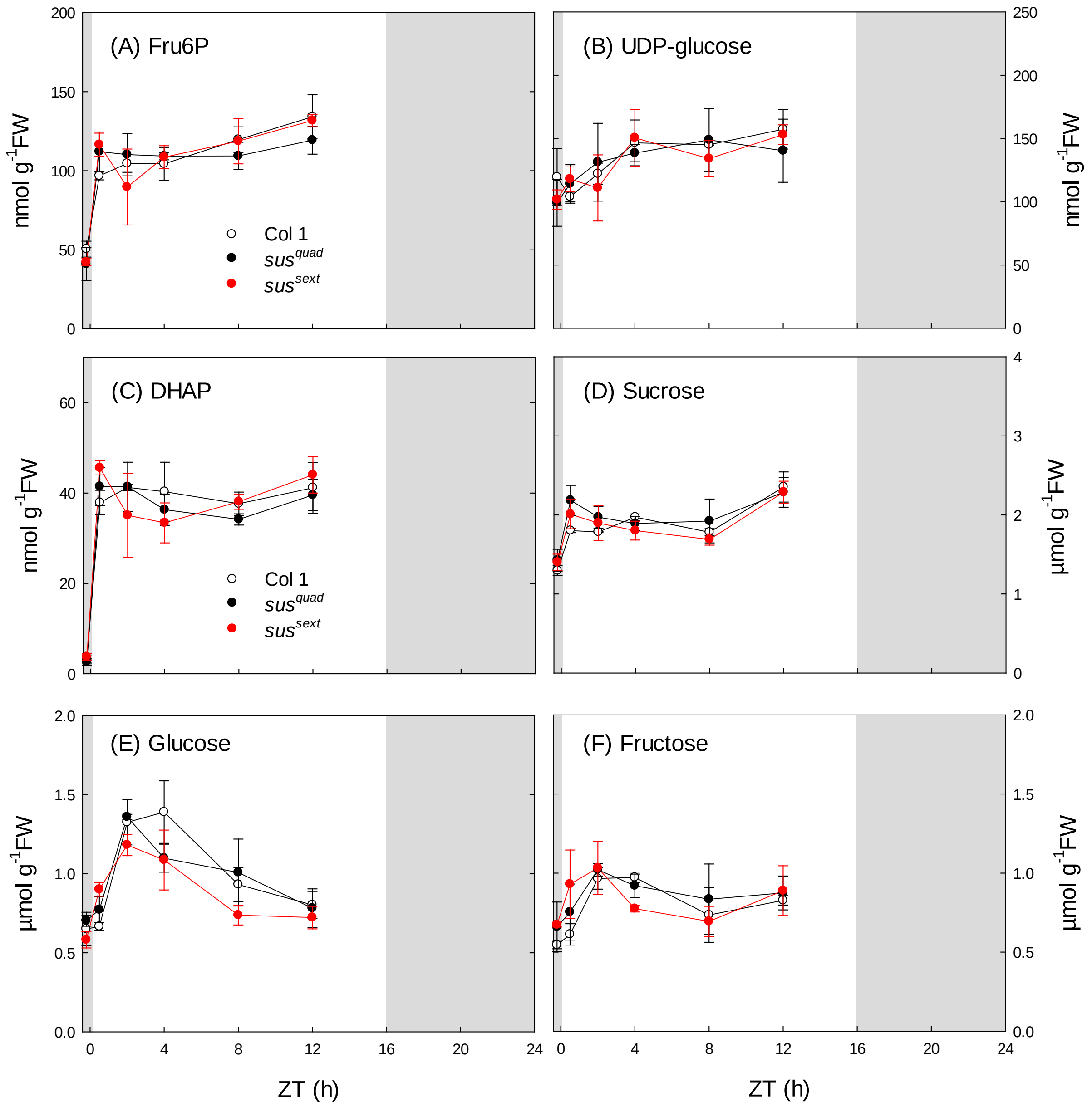


**Supplemental Figure S4. Metabolite levels in wild-type Arabidopsis and *sus* mutants.**

Wild-type Col-0, and the *sus1234* mutant (*sus*$^{quad}$) and *sus123456* (*sus*$^{sext}$*-1*) mutants were grown in long-day conditions (16 h photoperiod). At 25 days after germination, rosettes were harvested just before dawn (ZT-0.2) and at intervals from ZT0.5 to ZT12 for measurement of (A) fructose 6-phosphate (Fru6P), (B) UDP-glucose, (C) dihydroxyacetone-phosphate (DHAP), (D) sucrose, (E) glucose and (F) fructose. Data are from the same experiment as Fig. 5 and presented as mean ± S.D. (*n*=3). *P*-values for all genotype x genotype comparisons are shown in Supplemental Table S2.

**Supplemental Table S1. Statistical analysis of metabolite data in Figure 3.**

Metabolite levels in rosettes of Col-0, *sus1234* (*sus*$^{quad}$), *pgm* and *adg1* were compared across all sampling times by two-way ANOVA with Tukey's Honestly Significant Difference test. In columns 2-5, letters indicate significant differences between genotypes, with the *P*-value shown in column 6 and level of significance in column 7. *$P<0.05$; **$P<0.01$; ***$P<0.001$; n.s., not significant.

| **Metabolite** | **Col-0** | ***sus*$^{quad}$** | ***pgm*** | ***adg1*** | ***P*-value** | **Significance** |
|---|---|---|---|---|---|---|
| Glc6P | b | ab | a | a | <0.001 | *** |
| Glc1P | bc | ab | c | a | 0.166 | n.s. |
| ADP-Glc | a | a | b | b | <0.001 | *** |
| starch | a | a | b | b | <0.001 | *** |

**Supplemental Table S2. Statistical analysis of metabolite data in Figure 5 and Supplemental Figure S4.**

Metabolite levels in rosettes of Col-0, *sus1234* (*sus*$^{quad}$) and *sus123456* (*sus*$^{sext}$*-1*) were compared across all sampling times by two-way ANOVA with Tukey's Honestly Significant Difference test. In columns 2-4, letters indicate genotypes that were not significantly different for a given metabolite, with the *P*-value shown in column 5 and level of significance in column 6. n.s., not significant.

| Metabolite | Col-0 | *sus*$^{quad}$ | *sus*$^{sext}$*-1* | *P*-value | Significance |
|---|---|---|---|---|---|
| Glc6P | a | a | a | 0.518 | n.s. |
| Glc1P | a | a | a | 0.382 | n.s. |
| ADPG | a | a | a | 0.774 | n.s. |
| Starch | a | a | a | 0.228 | n.s. |
| Fru6P | a | a | a | 0.981 | n.s. |
| UDPG | a | a | a | 0.752 | n.s. |
| DHAP | a | a | a | 0.973 | n.s. |
| Sucrose | a | a | a | 0.360 | n.s. |
| Glucose | a | a | a | 0.535 | n.s. |
| Fructose | a | a | a | 0.471 | n.s. |

**Supplemental Table S3. Oligonucleotide primers for PCR genotyping.**

| **Primer** | **Sequence (5´→3´)** |
|---|---|
| *SUS genomic primers* | |
| SUS1-LP | ATGGCAAACGCTGAACGTATG |
| SUS1-RP | CTCAAGAGTGCAAGGATCAGG |
| SUS2-LP | ATGCGGAGACAAAATCACAAC |
| SUS2-RP | TAAGACTGTGAAAGTTGATGG |
| SUS3-LP | ATCGATGTGTTTGATCCGAAG |
| SUS3-RP | TTGGAGACCAGCGTCTGATAC |
| SUS4-LP | ATGGCAAACGCAGAACGTGTAA |
| SUS4-RP | CTTAGTCTTCTCCAAAGCATG |
| sus5-Fw | CGGATGACTCTATCTATTTCCCT |
| sus5-Rv | GAGACGTACATGTGTTCGTCATTC |
| sus6-Fw | ATGCTTGCTGTGATTGTTGTCTCG |
| sus6-Rv | CGAGGTAAGGGTAGATATCGAATC |
| *T-DNA left border primers* | |
| LBb1.3 | ATTTTGCCGATTTCGGAAC |
| P66 | CCCCTGCGCTGACAGCCGGAACACG |
